# Systematic evaluation of peptide property predictors with explainable AI technique SHAP

**DOI:** 10.1101/2025.11.19.689259

**Authors:** Joel Lapin, Maksim Iuzhaninov, Adrian Johannes Hölzlwimmer, Mathias Wilhelm

**Author notes:** Corresponding Authors Joel Lapin, School of Life Sciences, Maximus-von-Imhof-Forum 3, Technical University of Munich, Freising, DE. Mathias Wilhelm, School of Life Sciences, Maximus-von-Imhof-Forum 3, Technical University of Munich, Freising, DE.

## Abstract

Deep learning models are often characterized as black boxes because their layers of various mathematical transformations and activation functions are practically uninterpretable and have little meaning to the user. However, explainable AI methods exist to attribute model output predictions to its inputs. Shapley additive explanations (SHAP) is one such method that directly quantifies the inputs’ contributions and qualitatively addresses the question of why a model makes a particular prediction. SHAP generally is hampered by its computational cost, which scales very poorly with large inputs. Therefore peptide property predictors that take as input amino acid sequences, roughly between the lengths of 10-30 amino acids, represent ideal systems for applying SHAP. In applying SHAP to models that predict retention time, collisional cross section, peptide flyability, and fragment intensity, we obtain the relative influence that each amino acid has on predicting each property, and furthermore can rationalize the values from the perspective of the amino acids’ chemistry. Simply correlating the average shapley values per amino acid type over an entire validation dataset has yielded high correlation to published amino acid indices that are highly related to the property being predicted. For instance, our average shapley values for retention time had a 0.973 Pearson correlation with experimentally measured amino acid retention indices at pH 2. In applying SHAP on the Prosit fragment intensity prediction model, there is strong agreement with the mobile proton model, specifically demonstrating the effect of basic amino acids, the proline effect in charge-remote fragmentation, and positively charged amino acids in charge-directed fragmentation. We also use SHAP in a targeted experiment to demonstrate the *Pathways in competition* behavior of the model, and reveal a very discrete decision process based on basic residues and the fragment charge. While SHAP in this work was applied to models of well understood properties/systems, there is great potential to explain less studied areas of peptide chemistry to provide insights into their mechanisms.

## Introduction

Tandem mass spectrometry (MS2) has rapidly progressed to provide time and cost effective proteomics analyses, often utilized in order to identify and quantify the proteins present in a biological sample. In bottom up proteomics, single experiments will often produce millions of individual MS2 spectra, many of which contain no biologically relevant information, requiring the use of computationally intensive software tools to search the spectra. Standard practice is to search each spectrum against theoretical fragmentations of a library of peptide sequences expected to be present in the sample^1–5^.

The matching process between MS2 spectra and a sequence database involves obtaining candidate peptides by matching a precursor mass, and then proceeding to match the m/z’s of observed fragment peaks to theoretical fragments from the candidates^6,7^. The search space amongst candidate peptide sequences can be very large and inevitably leads to ambiguities, wherein many candidates’ theoretical spectra match the observed peaks reasonably well. The lack of specificity in matching can be greatly rectified using predicted intensities for the candidate peptides^8^, requiring a model of peptide fragmentation. It is well known that fragmentation amongst backbone amide bonds is most often non-uniform and highly sequence dependent, as well as being affected by the energy applied in the spectrometer under collision-induced dissociation (CID) and high energy collision-induced dissociation protocols (HCD)^9,10^. The uniqueness of MS2 spectra, intensities included, based on their sequence greatly increases the specificity of identification.

The understanding of the chemistry involved in fragmentation has long been incomplete, only having the ability to describe general trends for specific amino acids. The mobile proton is generally the accepted model that explains, qualitatively, certain amino acid’s relation to intense b- and y-ions, or the lack thereof^9–14^. For example, this theory rationalizes why peptides show enhanced fragmentation N-terminal to proline, or C terminal to glutamic and aspartic acid, or more generally in the vicinity of the most basic residues. Yet, the appearance of spectral intensities often cannot be rationalized beyond a limited number of qualitative observations, and “rules of thumb” for fragmentation^13^.

Machine learning, and specifically deep models have largely solved the problem of intensity prediction, as evident by their predictions’ strong agreement with experimentally obtained spectra^8,15–18^, and applications to significantly improve peptide identification^19–21^. These models have also been extended to other peptide property prediction problems, including but not limited to retention time, collisional cross section^22,23^, and peptide detectability^24^. Deep learning is quite unique in the sense that the model developer generally does not need an expert’s understanding of the underlying problem being modeled. With only a simple understanding that sequence, charge, and collision energy will affect fragmentation in varied ways, one could architect a deep neural network to connect these inputs to the ultimate output space, and yield a highly accurate model with enough training data. One presumes that a well trained model for intensity prediction has “learned” the complex chemokinetic rules of fragmentation as it relates to the amino acid sequence of the peptide being predicted.

Inspection of the model’s internal weights and calculated features, though, gives little insight into what “rules”, if any, the model has learned for peptide fragmentation. Deep learning models, in a literal sense, are not black boxes; each of their outputs can be written as a closed form expression that is a function of the model’s inputs. Yet the meaning of all the model’s operations and latent features, with respect to the problem being solved, are largely opaque, and practically impossible to interpret as to why the model makes its predictions. Methods for explainable AI exist, which can interrogate trained models’ behavior to reveal their modus operandi in non-abstract ways^25^.

Two related model-agnostic perturbation methods, Local Surrogate Models (LIME)^26^ and Shapley Additive Explanations (SHAP)^27^, generally are able to ascribe, quantitatively, importance to any inputs of choice. These methods work by exhaustively altering the original input to the model, gauging the effect on the output for every alteration made. For intensity prediction models, and more broadly peptide property predictors (PPP), this means mutating the amino acids of a peptide and quantifying the change in the model’s output(s). Of particular interest in this study is the SHAP method that explains the model through shapley values (SV)^28^, which have the desirable property of providing global explanations across an entire dataset. One previous study of note has served as a proof of concept for SHAP applied to biological sequences, wherein the authors calculated SV for peptide collisional cross section and binding affinity models^29^. Furthermore, SHAP is model-agnostic, i.e. it only requires a black box predictor to take inputs and produce outputs in order to be calculated, we can conveniently utilize the growing inventory of publicly available PPPs from the online prediction platform Koina^30^. This not only disencumbers the user of training their own models, but also obviates the need for GPU accelerators, which are often necessary for SHAP applied to deep learning models. This work potentially makes SHAP easily accessible to any model, past, present, or future, that uses amino acid sequences as the primary input.

Herein we apply SHAP to PPP models relevant to bioinformatics, namely intensity, retention time, collisional cross section, charge state, and time of flight prediction. We utilize SHAP not only to demonstrate that the model’s SV ascribed to the amino acids is consistent with known physicochemical attributes in the underlying predicted properties, but also to demonstrate the explanations’ potential to expand scientific knowledge for the peptide properties. By performing in depth analysis on the SV, we can highlight the subtleties of peptide fragmentation that have attracted less attention in the literature heretofore.

## Results

### Peptide Property Predictors: Retention Time, Collisional Cross Section, Peptide Flyability

We have generated explanations for a handful of models, all of which take peptides as inputs, but whose output space is small relative to the intensity model, often a single value. These models are the Prosit indexed retention time model (iRT)^8^, AlphaPeptDeep collisional cross section (CCS) model^18^, and Pfly, a recently published model that classifies peptide flyability^24^. In contrast to the intensity model, which takes additional inputs precursor charge and collision energy, all three of these peptide property predictors only take the peptide sequence as input.

In the liquid chromatography (LC) step of MS, peptide separation is critical to deconvoluting the analysis in MS2 (decreasing chimericity), and improving peptide identification. This can be measured by the peptides’ retention time, which is the rate at which the peptides elute in the column fed to the spectrometer. Retention time is ultimately a property of the chemical makeup of the peptide’s amino acids, and thus it can be modeled with reasonable accuracy^8,31^. We applied our SHAP calculation to the Prosit iRT model that predicts indexed retention time, accounting for the calibration between measurements from different machines.

As an initial proof of concept, we calculated the SV for Prosit’s iRT prediction over our entire dataset. Then, we took the mean SV for every amino acid and correlated them with a list of over 600 published normalized amino acid indices, of which each index contains 1 value for each of the 20 canonical amino acids^32^. As a demonstration, Figure 1 displays scatter plots with 2 example indices, retention coefficient at pH 2^33^ and transfer free energy to surface^34^, which are highly correlated and anticorrelated, respectively, with mean iRT SV. In this case, the former has a high positive correlation (0.921) and the latter has a high negative/anti correlation (−0.873).

**Figure 1:**
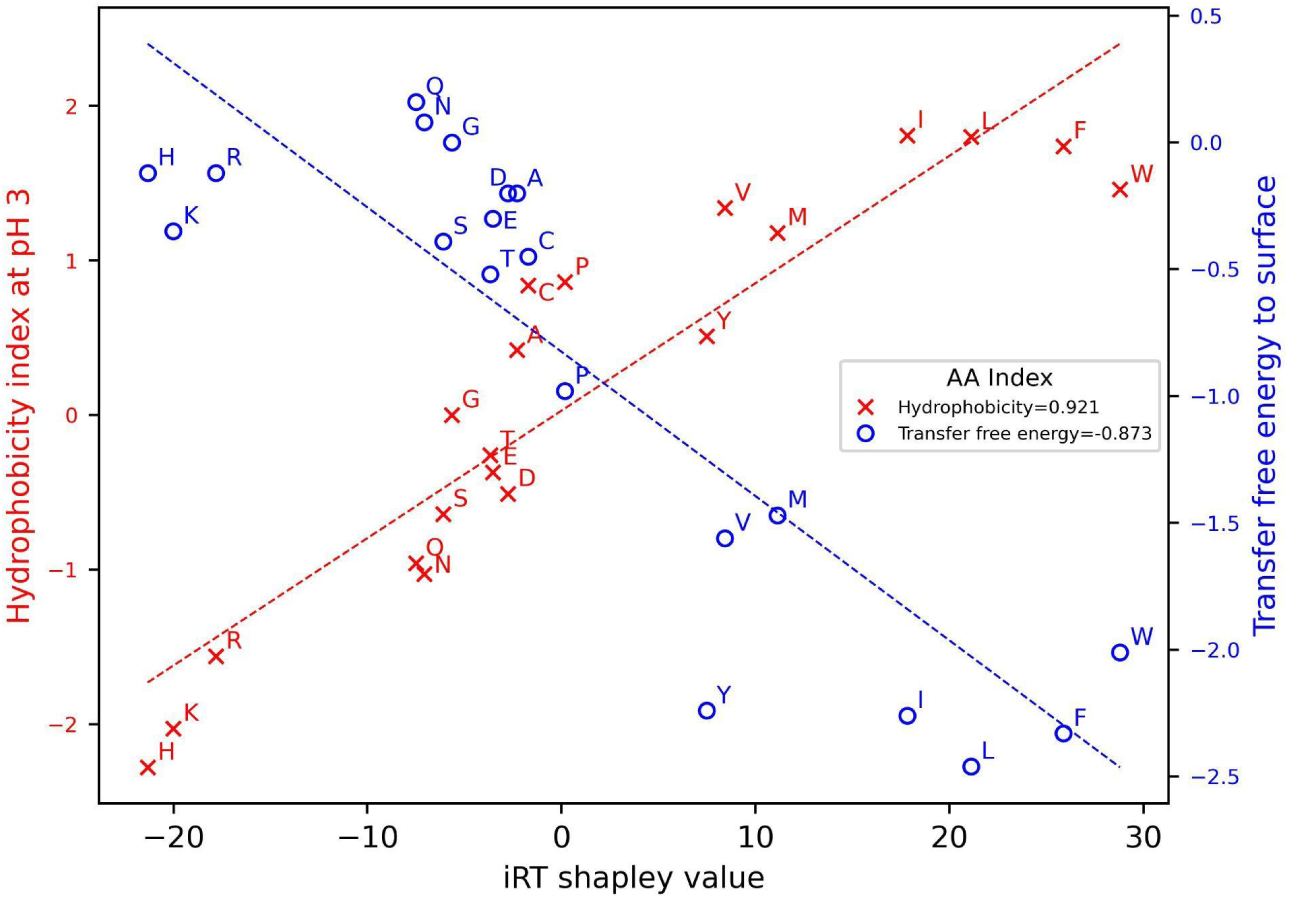
Scatter plots of average shapley values for the 20 canonical amino acids with experimentally determined amino acid indices *Hydrophobicity index at pH 3*^35^ and *Transfer free energy to surface*^34^. The legend displays the correlations between the shapley values and the indices; hydrophobicity is highly correlated and transfer free energy is highly anticorrelated. Each axis’ units are on the specific metric’s respective scale and are not directly comparable with each other.

Table 1 displays the three top absolute correlation/anticorrelations amongst all amino acid indices. The two top entries for retention time are studies where retention coefficients in LC for each amino acid are experimentally determined. For example, for *Retention coefficient at pH 2*^33^ each amino acid’s contribution to retention time is calculated by measuring the retention time for synthetic peptides in which two interior amino acids are synthesized to the amino acid of interest. The third top correlated index is related to hydrophobicity of amino acids, equivalently amino acid polarity, which has long been identified as a physicochemical property that is tightly linked to peptide retention^36^.

**Table 1:** Top absolute correlations for mean shapley values with Amino acid indices.

| Amino acid index | Correlation |
| --- | --- |
| Retention time |  |
| Retention coefficient at pH 2<sup>33</sup> | 0.973 |
| Retention coefficient in NaH<sub>2</sub>PO<sub>4</sub><sup>37</sup> | 0.932 |
| Hydrophobicity index at pH 3<sup>35</sup> | 0.921 |
| Collisional cross section |  |
| Helix termination parameter at position j+1<sup>38</sup> | -0.700 |
| RF value in high salt chromatography<sup>39</sup> | -0.699 |
| Weights for coil at window position of -6<sup>40</sup> | 0.696 |
| Peptide Flyability |  |
| Non-flyer: Information measure for middle turn<sup>41</sup> | 0.584 |
| Weak-flyer: Net charge<sup>42</sup> | -0.763 |
| Intermediate-flyer: Information measure for middle turn<sup>41</sup> | -0.642 |
| Strong flyer: Principal component IV<sup>43</sup> | -0.618 |

We then looked closer at the effect that position of an amino acid in a peptide may play in retention time by producing the heatmap in Figure 2A. The cells in this heatmap are now the mean SV for specific combinations of the amino acid and its relative position in the peptide. The basic amino acids which are likely to carry a charge, i.e. arginine, histidine, and lysine, have the strongest negative effects on retention time. Although the effect is not especially dramatic, the

**Figure 2:**
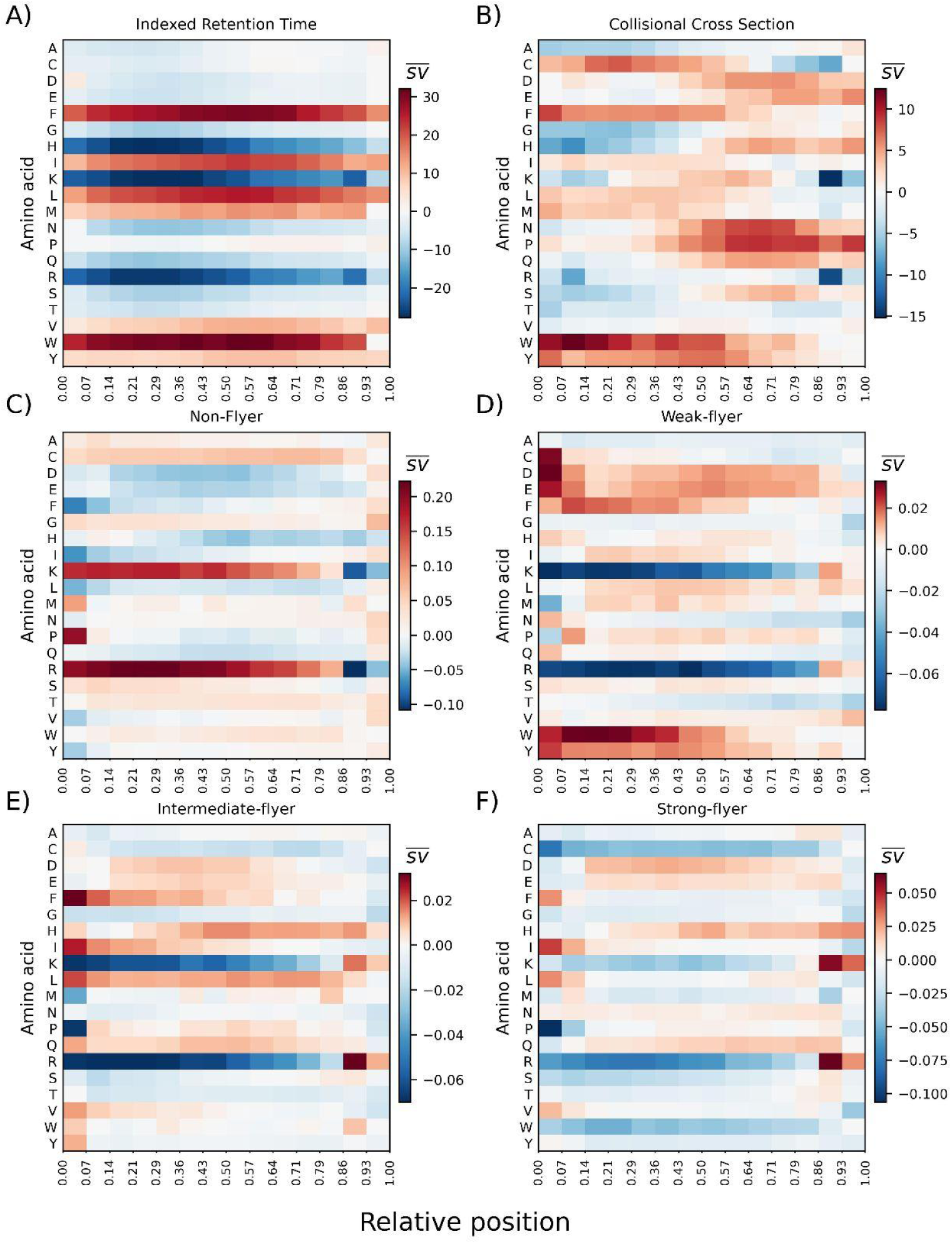
Amino acid-position mean shapley value heatmaps for 3 peptide property predictors: A) indexed retention time, B) collisional cross section, and C-F) flyability prediction for Non, Weak, Intermediate, and Strong-Flyer’s, respectively. Each entry represents the mean shapley value (s̅v̅) for the combination of amino acid and peptide position over our entire dataset. Position is represented by the binned fractional position of an amino acid from the N- (0) to the C-terminus (1). Absent/blank entries signify a lack of data available for a particular amino acid-position combination.

SV are larger towards the center of peptides, and slightly attenuated at the termini, especially the C-terminus. There are strong positive effects from phenylalanine and tryptophan, which both contain hydrophobic conjugated ring structures in their side chains. The same can be said, to a lesser extent, for leucine and isoleucine, which contain a simple hydrocarbon side chain. Again the effect of position is stronger when these amino acids lie in the middle of the peptide and less so on the termini. It should be noted that the model’s positive attribution for these amino acids is not necessarily reflective of the residues’ promotion of peptide mobility, but rather their absence of a retention effect on peptides in LC.

With the CCS model in Table 1, there are two indices that relate to peptide structure and conformation, specifically helix formation. An amino acid’s proclivity towards stable helix formation will give the peptide a more compact structure, and thus lower the collisional cross section. The next highest index, *RF value in high salt chromatography*^44^, which measures an amino acid’s relative hydrophobicity, has a strong anticorrelation. This is also related to peptide conformation, as hydrophobic amino acids will preferentially be buried from a solvent, encourage more compact structures, and lower CCS^45^.

While previous studies have applied SHAP to CCS models^29,45^, the effects of position seen in this study slightly differ in several subtle ways. In Figure 2B most amino acids show a polarity in mean SV from the N- to C-terminus. The amino acids that encourage higher CCS, i.e. proline and tryptophan, show opposite polarities; from the N- to C-terminus proline increases in SV, and tryptophan decreases. A similar observation can be made for the amino acids with more negative average SV, such as alanine, glycine, histidine, and serine; all four of these amino acids increase in value from the N- to C-terminus. The aforementioned study^29^ found an overall positive bias for amino acids at the termini. This applies to some amino acids for our calculations, but very much depends on the type of residue.

The final peptide predictor evaluated (besides intensity) predicts peptide flyability^24^. The model classifies peptides amongst four classes: 1) Non-flyer, 2) Weak-flyer, 3) Intermediate-flyer, and 4) Strong-flyer. This attribute is a relative measure of a peptide’s detectability, or likelihood of producing detectable intensity in MS1. We expect correlations to be generally lower than iRT or CCS because detectability in a spectrometer is the aggregate of many properties, e.g. the efficiency of the digest, LC phase mobility, whether it is ionized, etc. There are generalizations, though, that predispose peptides to detectability; for instance more hydrophobic amino acids will be better retained in the LC phase of MS1, increasing the likelihood of the peptide’s detection. Each of the four classes’ top correlation is displayed in Table 1. There is a recurrence of attributes relevant to peptide helicity. *Information measure for middle turn*^41^ is a measure of an amino acid’s proclivity towards coiling; it is therefore consistent that its correlation flips from positive to negative going from non-flyers to intermediate-flyers. The *Principal Component IV*^43^ index is also an amino acid’s measure of helicity, albeit in a more indirect way. The values of this attribute from a principal component analysis of protein properties are related to an amino acid’s propensity to form hydrogen bonds, which are necessary between backbone amine and carbonyl groups in order to form alpha helices^43^. The anticorrelation with class Strong-flyer’s SV implies the counterproductive role that helicity plays in peptide detection. Finally Weak-flyers have a strong anticorrelation with amino acid net charge, reflecting the basic amino acids having a largely positive effect on flyability^42^.

### HCD Intensity Model: Comparison to the Mobile Proton Model

Our analysis then proceeded to the Prosit intensity prediction model, whose 174 ion output space requires a more detailed analysis than the PPP models. A comprehensive model/framework that has served to qualitatively rationalize fragmentation spectra is the mobile proton model. The theory is largely developed on the results of targeted studies of various peptides^12,14,46,47^, electronic structure calculations to understand the intermediate chemical species involved^48,49^, and statistical analyses^50,51^. Charged peptides, i.e. with extra protons beyond the neutral molecule (perhaps originating on a basic residue’s side chain), can have mobile protons migrating along the molecule upon excitation, triggering cleavages of amide bonds via charge-directed fragmentation. When a proton becomes mobile it will find otherwise less favored protonation sites, creating intermediate species that lead to backbone cleavages. Certain amino acids, as a result of their side chain chemistry, can enhance or suppress the likelihood of fragmentation at their corresponding amide bonds^50^, e.g. basic residues, such as arginine, histidine, and lysine, are capable of sequestering protons, preventing them from becoming mobile and thus requiring more energy for excitation. In contrast to charge-directed fragmentation, charge remote fragmentation occurs when charged side chains, such as aspartic or glutamic acid, are involved in nucleophilic attack of the backbone, triggering the fragmentation event. Ultimately charge-directed and charge-remote fragmentation pathways are in competition with each other, with the more kinetically favored pathways increasing the intensity of their resultant fragment ions^52^.

One of the most common rules that is well described by the mobile-proton model is the enhanced cleavage of an amide bond, N-terminal to proline (Xaa-Pro), to form intense y-ions^9,11,12,14,46,50,53^. Proline is a special amino acid, having a side chain that connects to its backbone nitrogen, forming a pyrrolidine ring. This renders proline’s nitrogen more basic than the amino groups of other amino acids and therefore more likely to hold the mobile proton which initiates fragmentation^46^. In Figure 3D-F there are different length y-ions, for which proline in position 0, i.e. on the N-terminus of the y-fragment, overwhelmingly makes highly positive contributions to the predicted intensity. Alternatively, Proline is said to disfavor fragmentation when lying on the C-terminal side of the fragmentation point, which is evident by the low SV for P at the −1 position of y-ions. This is explained by the Proline pyrrolidine ring sterically hindering the formation of a ring between adjacent backbone carbonyl groups, a necessary intermediate species for C-terminal fragmentation^9^. For b-ions (Figure 3A-C) there is the exact opposite effect compared to y-ions; proline makes very negative contributions to intensity at the 0 position (N-terminal fragmentation), and positive contributions at the −1 position (C-terminal fragmentation).

**Figure 3:**
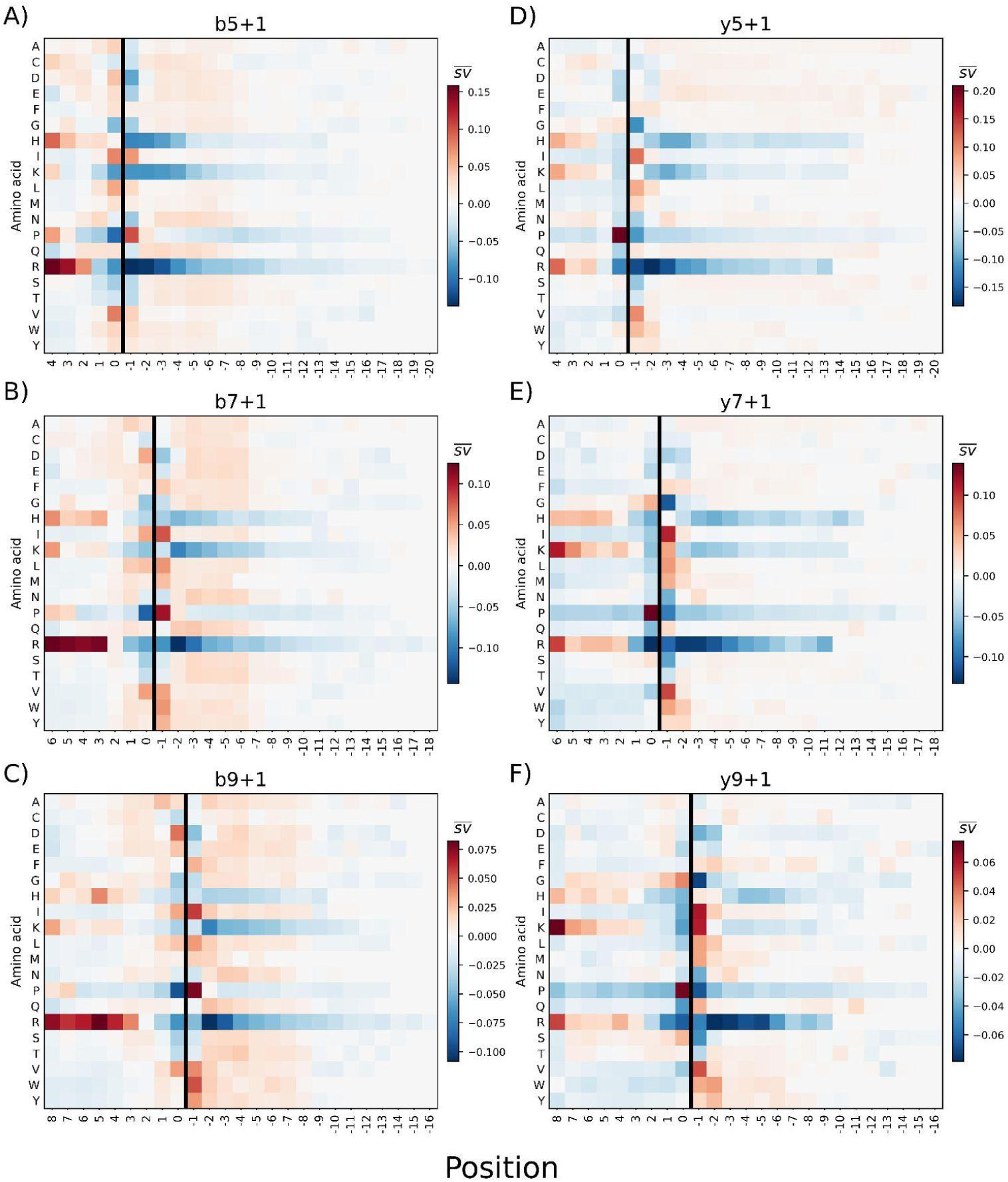
Heatmaps of mean shapley values (s̅v̅) for all combinations of amino acids and positions, for b-(left) and y-ions (right) of length 5 (top), 7 (middle), and 9 (bottom). The mean values are taken over the entire peptide test set. The coordinates of position in the x-axis are numbered relative to the ion fragment point, denoted by a solid black line, where 0 is the last amino acid in the fragment (the fragment’s N-terminus for a y-ion, C-terminus for a b-ion). Positive numbers increase towards the N-terminus for b-ions and towards the C-terminus for y-ions, and negative numbers decrease (larger negative value) towards the respective opposite termini.

The tendency of amino acids with acidic side chains, which includes aspartic acid (Asp-X) and glutamic acid (Glu-X), to produce intense b-ions by fragmentation on their C-terminus is well documented and also rationalized through the mobile proton model^9,11,13^. For these amino acids the proton that initiates fragmentation comes from the acidic side chains in a charge-remote fragmentation. The global mean SV of the b-ions (Figure 3A-C) confirm this observation, where both aspartic acid and glutamic acid have a light positive effect (the former being stronger) when N-terminal to the fragment point.

Overall there is strong agreement between the global mean SV of Prosit with the hypotheses of the mobile proton model. More specifically, our SHAP calculations confirm the trends seen for both charge-directed fragmentation, e.g. the proline rule for y-ions, and charge-remote fragmentation by acidic amino acids. It should be cautioned that the heatmaps of Figure 3 are mean SV over all explained peptides, and that the combination of amino acid and position in the sequence is not the sole determinant of the contribution to fragment intensity. In fact there are a range of SV for any amino acid-position combination from our dataset, whose value is conditioned on surrounding amino acids as well as peptide length and charge (see *Pathways in competition model*).

### Aliphatic Amino Acid Side Chains

From our calculations on HCD spectra, a consistent pattern is the promotion of both b-and y-ion intensity by amino acids leucine, isoleucine, and valine. These amino acids with simple hydrophobic aliphatic side chains exert their effect when just outside of the fragment, at the −1 position in Figure 3 heatmaps; specifically this is C-terminal to b-ions and N-terminal y-ions. The enhanced C-terminal cleavage of these three amino acids has been previously documented in analyses of observed intensity patterns in spectra^50,53^, and is suggested to occur because of the side chains’ restricted rotational conformations creating reactive conformations for fragmentation^54^.

Previous work analyzing fragmentation intensities did not find enhanced N-terminal cleavage, though, for these amino acids^50^. In Figure 4A,C isoleucine, leucine, and valine at the −1 position are all amongst the top 20 tokens in terms of positive SV. This is especially true for y5+1 (Figure 4C), where I-^-^1, V-^-^1, and L-^-^1 are all in the top 6 by mean SV. Further evidence of aliphatic amino acids increasing fragmentation are in the bi-Token plots in Figure 4B,D for pairs of amino acids on either side of the fragment point (−1 and 0). Ion b5+1 has I, L, or V on either side of the fragment point in 13 of its top 20 bi-tokens (Figure 4B), and 11 of the top 20 for y5+1 (Figure 4D, specifically at position −1). It has been previously shown that these 3 aliphatic amino acids promote N-terminal cleavage of proline, with valine>histidine>aspartic acid>isoleucine>leucine^53^. It should be noted that the statistics in the Figure 4A,C boxplots are taken over all occurrences of I,L,V at specific positions in our dataset, and not only when N-terminal to proline. It can be inferred from our results that the model treats I,L,V as facilitators of cleavage on both termini, with isoleucine and valine often having greater effect than leucine.

**Figure 4.**
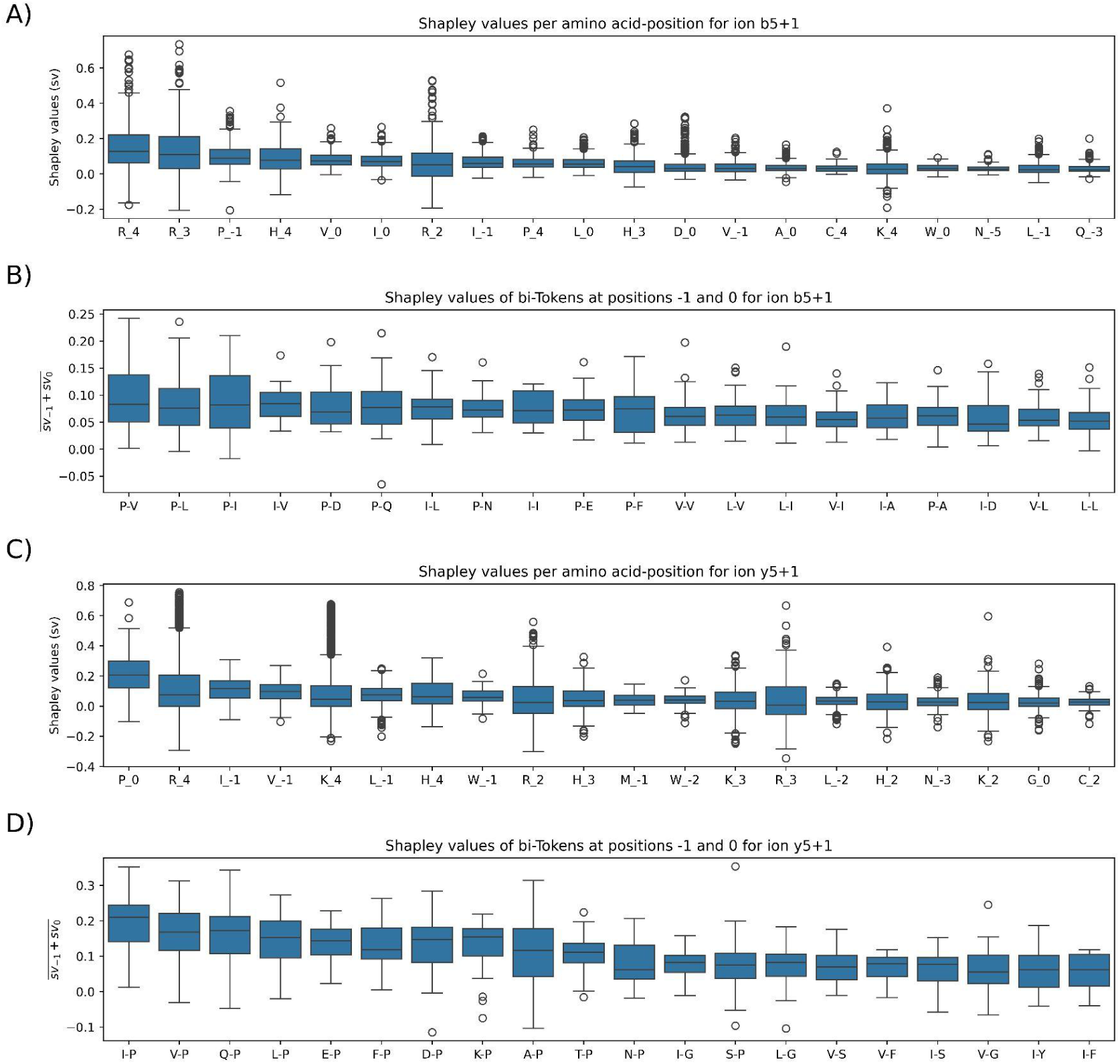
Boxplots of top 20 major tokens, sorted high to low by highest mean SV. In A,C) the tokens are unique amino acid - position combinations for b5+1 and y5+1, respectively, and in B,D) the tokens are combinations of 2 amino acids at the −1 and 0 positions for the same 2 ions, respectively. The coordinate system follows our convention where the C- and N-termini of b- and y-ion fragments, respectively, are the 0 position; positive increasing numbers are amino acids contained in the fragment towards the other terminus, and negative numbers are outside of the fragment towards the opposite peptide terminus. For bi-Token plots, amino acids for every combination are listed in position order −1-0.

### Pathways in Competition Model - A Targeted Experiment

While the heatmaps of mean SV in Figure 3 are useful for giving an overview, it is an oversimplification of the model’s complex behavior to reduce each amino acid-position to a single number. The boxplots of Figure 4A,C show that amino acids can take on a wide range of SV at a specific position. This indicates that the contributions of amino acids have further conditions that determine their effect on the intensity. We elucidated one of these conditions, specifically for basic amino acid arginine, by explaining custom designed peptides that are altered in targeted ways.

The pathways in competition model describes the competition of charge-directed and charge-remote fragmentation, for which basic amino acids play the central role. The effect of acidic amino acids on fragmentation is amplified in peptides with more internal basic amino acids, arginine, histidine, or lysine. In the pathways in competition model, this observation is rationalized by basic amino acids’ ability to hold and strongly sequester a proton, lowering its mobility and capacity to initiate charge-directed fragmentation^52^. That is, as the energy required for charge-directed fragmentation pathways (e.g. proline effect) is raised by basic amino acids, charge-remote pathways (e.g. C-terminal fragmentation to aspartic acid) become more dominant. This effect is reported in a statistical study of 5500 HCD spectra, where peptides characterized as partially or non-mobile (based on the precursor charge and basic amino acid content) show enhanced C-terminal cleavage of aspartic acid residues and suppressed N-terminal cleavage of proline^50^.

To probe this effect in the intensity model, we observed the SV of peptides with varying numbers of arginines in specific locations along the peptide. The starting point is a largely poly-alanine peptide, 9 residues in length, precursor charge +2, with an aspartic acid and a proline in the 3rd and 6th positions from the N-terminus, respectively. In this setting, the aspartic acid favoring charge remote fragmentation on its C-terminal side competes with the proline favoring charge directed fragmentation on its N-terminal side. This peptide was altered by adding one arginine in three different locations, and then a second arginine in seven unique configurations. The alanines are expected to often have low contributions, as observed in Figure 3.

Table 2 shows the SV of aspartic acid, proline, and arginines for 3 ions that fragment on the C-terminal side of aspartic acid; Table 3 shows the same for 3 ions that fragment on the N-terminal side of proline. For the base sequence without a single arginine residue (AADAAAPAA), the b3+1 and y3+1 are the most intense ions. As one arginine is added at the C-terminus (AADAAAPA**R**), making the peptide tryptic, y6+1 now becomes an intense ion, owing to a large contribution from the tryptic arginine (it is notable that the same arginine has a slightly negative contribution to the y3+1 ion). When the arginine is moved to the other terminus (**R**ADAAAPAA), in a similar respect to the y-ions with a C-terminal arginine, the N-terminal arginine now has a very positive SV in b6+1 and slightly negative in b3+1; y3+1 still has considerable intensity due to the proline, but is somewhat attenuated by the N-terminal arginine. When a second arginine is substituted for another alanine, the peptides are now classified as non-mobile since both precursor charges can be sequestered by the basic residues. The mobile proton theory accordingly predicts that this should energetically discourage charge-directed fragmentation, elevating the predominance of charge-remote fragmentation, i.e. C-terminal to aspartic acid. Among the 7 non-mobile peptides, the C-terminal cleavage ions contain the base peak in 5 cases, mostly for ion y6+2. This is evident in all peptides where y6+2 contains both arginines (AADAAAP**RR**, AADA**R**APA**R**, AADA**RR**PAA, AAD**R**A**R**PAA); the 2 arginines presumably hold the 2 charges that give the ion its +2 fragment charge. Only when y6+2 does not contain both arginines (**RR**DAAAPAA and **R**ADAAAPA**R**), its intensity is 0. In both cases the arginine(s) outside of the ion have negative SV, revealing a very discrete behavior in the model. For **R**ADAAAPA**R**, y6+1 becomes the base peak, which still reflects how the model favors charge-remote fragmentation under non-mobile conditions.

**Table 2:** Shapley values of single peptides for 3 ions C-terminal to aspartic acid. All peptides have a collision energy of 30 and a precursor charge of +2; peptides are classified as *Mobile* if the number of arginines is less than 2. The model’s output intensity *f(x)* is shown beside the background dataset’s average intensity *E[f(x)]* for each ion. 4 ions’ intensity is displayed, including aspartic acid, proline, and the variable number of arginines, whose position varies as well. If there are 2 arginines, we differentiate them as one closer to the N-terminus and the other closer to the C-terminus. Red font denotes arginine inside of the fragment ion, blue font is outside.

| Sequence | Mobile? | b3+1 |  |  |  |  |  | y6+1 |  |  |  |  |  | y6+2 |  |  |  |  |  |
| --- | --- | --- | --- | --- | --- | --- | --- | --- | --- | --- | --- | --- | --- | --- | --- | --- | --- | --- | --- |
| | | $f(x)$ | $E[f(x)]$ | $D_0$ | $P_{-4}$ | $R_N$ | $R_C$ | $f(x)$ | $E[f(x)]$ | $D_{-1}$ | $P_3$ | $R_N$ | $R_C$ | $f(x)$ | $E[f(x)]$ | $D_{-1}$ | $P_3$ | $R_N$ | $R_C$ |
| AADAAAPAA | TRUE | 0.44 | 0.15 | 0.08 | -0.05 |  |  | 0 | 0. | -0.04 | -0.07 |  |  | 0 | 0.06 | 0 | -0.01 |  |  |
| AADAAAPAR | TRUE | 0.61 | 0.15 | 0.13 | 0 |  | 0 | 0.65 | 0.25 | -0.11 | -0.02 |  | 0.35 | 0 | 0.06 | 0.02 | -0.02 |  | 0.05 |
| RADAAAPAA | TRUE | 0.02 | 0.15 | 0.07 | -0.04 | -0.15 |  | 0 | 0.25 | 0 | -0.05 | -0.06 |  | 0 | 0.06 | 0 | -0.01 | -0.01 |  |
| AADARAPAA | TRUE | 0.09 | 0.15 | 0.04 | -0.01 | -0.19 |  | 0.41 | 0.25 | -0.08 | -0.01 | 0.26 |  | 0 | 0.06 | 0.04 | -0.03 | 0.06 |  |
| AADAAAPRR | FALSE | 0.25 | 0.15 | -0.03 | -0.03 | -0.05 | -0.04 | 0.04 | 0.25 | -0.13 | 0 | 0 | -0.07 | 1 | 0.06 | 0.23 | 0 | 0.36 | 0.34 |
| RRDAAAPAA | FALSE | 0.11 | 0.15 | 0.09 | -0.04 | 0 | -0.04 | 0.10 | 0.25 | 0.04 | -0.06 | 0 | 0 | 0 | 0.06 | 0 | 0 | -0.01 | -0.01 |
|  |  | f(x) | E[f(x)] | D<sub>0</sub> | P<sub>-4</sub> | R<sub>N</sub> | R<sub>C</sub> | f(x) | E[f(x)] | D<sub>-1</sub> | P<sub>3</sub> | R<sub>N</sub> | R<sub>C</sub> | f(x) | E[f(x)] | D<sub>-1</sub> | P<sub>3</sub> | R<sub>N</sub> | R<sub>C</sub> |
| AADAAAPAA | TRUE | 0.44 | 0.15 | 0.08 | -0.05 |  |  | 0 | 0. | -0.04 | -0.07 |  |  | 0 | 0.06 | 0 | -0.01 |  |  |
| AADAAAPAR | TRUE | 0.61 | 0.15 | 0.13 | 0 |  | 0 | 0.65 | 0.25 | -0.11 | -0.02 |  | 0.35 | 0 | 0.06 | 0.02 | -0.02 |  | 0.05 |
| RADAAAPAA | TRUE | 0.02 | 0.15 | 0.07 | -0.04 | -0.15 |  | 0 | 0.25 | 0 | -0.05 | -0.06 |  | 0 | 0.06 | 0 | -0.01 | -0.01 |  |
| AADARAPAR | FALSE | 0.04 | 0.15 | 0 | 0 | -0.19 | -0.02 | 0 | 0.25 | -0.05 | 0 | -0.14 | -0.06 | 1 | 0.06 | 0.29 | 0 | 0.33 | 0.32 |
| AADARRPAA | FALSE | 0.03 | 0.15 | -0.02 | 0 | -0.13 | -0.05 | 0 | 0.25 | -0.05 | -0.01 | -0.12 | -0.05 | 1 | 0.06 | 0.25 | 0 | 0.34 | 0.39 |
| AADRARPAA | FALSE | 0.03 | 0.15 | 0.03 | -0.02 | -0.15 | -0.02 | 0.03 | 0.25 | -0.04 | -0.05 | -0.28 | 0.16 | 1 | 0.06 | 0.31 | 0.01 | 0.24 | 0.36 |
| AADRRAPAA | FALSE | 0.07 | 0.15 | 0.03 | 0 | -0.09 | -0.04 | 0.21 | 0.25 | 0 | 0 | -0.18 | 0.16 | 0.88 | 0.06 | 0.27 | 0 | 0.22 | 0.30 |
| RADAAAPAR | FALSE | 0.84 | 0.15 | 0.29 | 0 | 0.10 | 0.21 | 1 | 0.25 | 0.18 | 0 | 0.02 | 0.44 | 0 | 0.06 | 0.01 | -0.02 | -0.03 | 0.03 |

**Table 3:** Shapley values for 3 ions N-terminal to proline. Same symbol representations as for. **Table 2**.

| Sequence | Mobile? | b6+1 |  |  |  |  |  | b6+2 |  |  |  |  |  | y3+1 |  |  |  |  |  |
| --- | --- | --- | --- | --- | --- | --- | --- | --- | --- | --- | --- | --- | --- | --- | --- | --- | --- | --- | --- |
|  |  | f(x) | E[f(x)] | D<sub>3</sub> | P<sub>-1</sub> | R<sub>N</sub> | R<sub>C</sub> | f(x) | E[f(x)] | D<sub>3</sub> | P<sub>-1</sub> | R<sub>N</sub> | R<sub>C</sub> | f(x) | E[f(x)] | D<sub>-4</sub> | P<sub>0</sub> | R<sub>N</sub> | R<sub>C</sub> |
| AADAAAPAA | TRUE | 0.32 | 0.097 | -0.01 | 0.09 |  |  | 0.01 | 0.03 | -0.01 | 0 |  |  | 0.68 | 0.115 | 0 | 0.49 |  |  |
| AADAAAPAR | TRUE | 0 | 0.10 | 0 | 0.06 |  | -0.19 | 0 | 0.03 | 0 | 0 |  | -0.02 | 0.47 | 0.12 | 0.04 | 0.32 |  | -0.07 |
| RADAAAPAA | TRUE | 1 | 0.10 | 0.01 | 0.2 | 0.38 |  | 0.03 | 0.03 | -0.02 | 0 | 0.06 |  | 0.22 | 0.12 | 0 | 0.35 | -0.20 |  |
| AADARAPAA | TRUE | 0.18 | 0.10 | -0.02 | 0.11 | -0.13 |  | 0.02 | 0.03 | -0.01 | 0.01 | -0.03 |  | 0.17 | 0.12 | -0.05 | 0.30 | -0.17 |  |
| AADAAAPRR | FALSE | 0.02 | 0.10 | 0 | 0.07 | -0.08 | -0.09 | 0 | 0.03 | 0 | 0 | -0.01 | -0.01 | 0.13 | 0.12 | -0.03 | 0.27 | -0.16 | -0.08 |
| RRDAAAPAA | FALSE | 0.58 | 0.10 | 0.03 | 0.27 | 0 | -0.05 | 1 | 0.03 | 0.01 | 0.09 | 0.38 | 0.39 | 0.29 | 0.12 | 0 | 0.33 | -0.02 | -0.04 |
| AADARAPAR | FALSE | 0.03 | 0.10 | -0.02 | 0.09 | -0.01 | -0.14 | 0 | 0.03 | -0.01 | 0.01 | 0.01 | -0.03 | 0.05 | 0.12 | -0.05 | 0.22 | -0.17 | -0.04 |
| AADARRPAA | FALSE | 0.04 | 0.10 | -0.13 | 0.09 | 0.01 | -0.15 | 0.05 | 0.03 | -0.07 | 0.02 | 0.04 | 0.01 | 0.07 | 0.12 | -0.18 | 0.23 | -0.05 | -0.05 |
| AADRARPAA | FALSE | 0.49 | 0.10 | -0.05 | 0.22 | 0.26 | -0.27 | 0.75 | 0.03 | -0.03 | 0.08 | 0.28 | 0.14 | 0.41 | 0.12 | -0.04 | 0.30 | 0.03 | -0.03 |
| AADRRAPAA | FALSE | 0.33 | 0.10 | 0.01 | 0.20 | 0.12 | -0.32 | 1 | 0.03 | -0.04 | 0.2 | 0.28 | 0.23 | 0.34 | 0.12 | -0.04 | 0.31 | -0.03 | -0.07 |
| RADAAAPAR | FALSE | 0.84 | 0.10 | 0 | 0.20 | 0.46 | -0.18 | 0.03 | 0.03 | -0.01 | 0.01 | 0.06 | -0.04 | 0.13 | 0.12 | 0.02 | 0.24 | -0.13 | -0.05 |

Arginine tends to make positive contributions when contained in an ion whose fragment charge is equal to the total number of arginines in the ion. When the number of arginines exceeds the fragment charge, their contribution becomes negative, especially for the R closer to the ion’s fragmentation point. For the 3 peptides where the two R’s are between D and P (AAD**RR**APAA, AAD**R**A**R**PAA, AADA**RR**PAA), at least one arginine (likely the one closer to the fragmentation point) makes a large negative contribution for y6+1 and b6+1. By contrast the contributions are large and positive for the charge +2 ions b6+2 and y6+2. This conditional behavior contrasts greatly with proline, which tended to make the same contribution for every ion in spite of the peptide’s changing chemical makeup. For example, proline largely had SV between 0.2-0.3 for y3+ of all peptides. These observations suggest that the model’s intensity prediction is largely driven by the basic amino acids in a very discrete manner, specifically their positions and amount in relation to the ion’s fragment charge and the peptide’s precursor charge.

Although the accuracy of these predictions cannot be verified since we do not have ground truth spectra for these peptides, we can still intuit that the model is sensitive to the mobile/non-mobile characterization of peptides. The results are consistent with the mobile proton and pathways in competition models of peptide fragmentation, insofar that the switch from a mobile to non-mobile peptide suppresses charge-directed fragmentation N-terminal to proline, and promotes charge-remote fragmentation C-terminal to aspartic acid.

## Discussion

We have demonstrated that SHAP produces explanations very faithful to the quantitative and qualitative “rules” known to govern the peptide properties relevant to mass spectrometry. The models used in this study represent an ideal application for SHAP, which scales very poorly with sequence size. By consolidating the input into its 10-30 individual amino acids, the peptides are short enough to produce accurate and reproducible explanations of the predictions. The peptide properties, i.e. iRT, CCS, detectability prediction, and HCD intensity spectra are well studied systems; there exists a history of background research and literature that we can use to rationalize the trends seen in experimental data. In running SHAP on all these models, we were largely able to confirm what would be expected given the background knowledge. For instance, average SV for the iRT model were highly correlated with retention indices from previous studies, and the HCD model was highly consistent with the mobile proton and pathways in competition models that rationalize trends seen in fragmentation spectra. Explanations in terms of the amino acid inputs have further use than simply confirming experimental observations, as they provide very granular data on the effects of different amino acids on the output.

Another important application is using SHAP to unveil the behavior and modus operandi of the model. It is known that deep learning models are desirable for their capacity to exhibit very complex behaviors and decisions. Prosit predicts continuous intensities for its fragment ions, but the model’s behavior can be very discrete depending on the fragment ion’s charge and the number of basic amino acids it contains. In Tables 2 and 3, a charge +2 fragment ion gets little predicted intensity if it contains only one arginine, but upon the addition of a second arginine there is suddenly a very large predicted intensity and the arginines’ SV go up considerably. This is a specific case in which the mean SV for an amino acid-position combination over the entire evaluation set is insufficient to explain the model’s behavior. For instance, whereas proline on the N-terminus of a y-ion always had high SV from the model, the basic amino acids’ SV rely on the configuration of basic amino acids in the fragment overall. To discern this behavior of the model from the predicted intensities alone would have been difficult to diagnose without the SV.

The focus of this study was to assess the potential for explainable AI, applied to peptide property predictors, to provide insights into the physical aspects of mass spectrometry. We applied SHAP to models for which we have a decent expectation of their predicted properties, i.e. what amino acids drive their predictions, and saw close agreement with both quantitative indices and qualitative observations in literature. For future work, we would like to apply SHAP to lesser known systems lacking theoretical models, such as the effects of post-translational modifications on spectra. We would also like to apply SHAP to lesser known fragmentation methods, e.g. ETD, ECD, UVPD, etc., whose mechanisms of fragmentation lack a comprehensive model like the mobile proton hypothesis. Our methods have potential to inform the development of those techniques, both in understanding the hardware as well as their application.

## Methods

### Data

All SV for the PPP models and the intensity HCD model were calculated on a dataset of 8,317 unmodified peptides originating from the ProteomeTools project^55^. In principle we could construct a dataset on any list of peptides which can be accepted as input by the model, although we elected to stick to only experimentally observed sequences (with the exception of the peptides in Tables 2 and 3). Peptides range in lengths from 6-30 amino acids, precursor charges +1-6, and normalized collision energies 20-35. Input to the model must be curated as a parquet file which contains a column that represents the SHAP input as a linear string array of all inputs the model needs to predict the peptide, i.e. peptide sequence, charge, and collision energy. See below (*SHAP*) in methods for more details. We provide the original input for this project in our Github repository for the project (https://github.com/jlapin1/shap-prosit).

### Models

Trained intensity, indexed retention time, CCS, and peptide flyability prediction models were sourced via the Koina public server^30^. This entails peptide inputs from the SHAP calculation being sent to the server and prediction values returned as pandas dataframes. The intensity model used was Prosit HCD 2019^8^, an accurate and widely implemented recurrent neural network, which produces 174 output ion intensity predictions. Since full b- and y-ion series are returned by the Koina server, the ion(s) of interest for the SHAP explanation must be indexed and extracted. The iRT model is Prosit 2019 iRT^8^, which is the same model architecture trained for index retention time. Retention time is only a single float returned by the server. The CCS predictor used is the AlphaPeptDeep CCS model^18^, which uses a composite architecture consisting of both recurrent neural network layers and convolutional layers. The peptide flyability predictor Pfly is a recurrent neural network, with an encoder/decoder framework, and outputs 4 classes of flyability probabilities.

### SHAP

Calculating SV via SHAP requires python’s *shap* library; all other calculation operations are carried out using the Numpy and Pandas libraries. The process begins with the dataset, which we organize in a parquet format. For each data instance, SHAP takes as input a linear array containing all input data necessary for model inference. For example, the intensity model data instances are the amino acid sequence, collision energy, and charge. The amino acid sequence is represented as strings for the amino acids, the collision energy as a float, and an integer for charge. The amino acid sequence is padded to its predefined max length with empty strings.

A model/predictor object must be specified. Any model must contain a function that takes the input string list and returns a float(s) for the prediction. Since the calculation of SV using SHAP is model-agnostic, accommodating any model requires writing an input hook to transform the input string into the appropriate input format for the backend model being used. For Koina, the amino acid strings, charge and energy must be collected into a python dictionary before submitting to Koina. Koina returns the predictions as a dataframe, from which the desired output is extracted for the SHAP calculation. If a local model for a deep learning framework such as Pytorch is used, the amino acid strings must be tokenized into integers, converted into torch tensors along with charge and energy, and then fed into the model.

A background dataset must be chosen at the beginning of the process and held constant for all calculations. Originally this is just a random subsampling of 100 peptides from our dataset. We then saved these peptides and thereafter reused them for all calculations to ensure consistency. From the background dataset the expectation value is calculated for the property being explained, e.g. the average intensity for the b5 ion, or the average retention time, over all background data instances. The python *shap* library’s backend for the calculation of SV requires that the input is the form of a binary vector. The binary vector indicates to the calculator what inputs to include in coalitions and what to exclude. For our calculations, the possible coalition members were always the amino acids up to the length of the actual peptide, and excludes padded amino acids, charge and collision energy, in effect leaving their values constant throughout the calculation. In principle charge and energy can be included in the explanation, but since they are precursor level information, in contrast to sequence level information, they would complicate analysis and thus were omitted.

The *shap* library generates binary coalition vectors, from which we create the altered peptide sequences. When a coalition vector contains a one, we use the original amino acid in that position from the peptide being explained. If the coalition vector contains a zero, then we substitute amino acids from the background dataset at positions equivalent to the zero. One complication of this method occurs when a peptide sequence from the background dataset is shorter than peptide sequence being explained, such that there are possible substitutions that result in a null/absent/pad amino acid at a position shy of the C-terminus, leaving null tokens within the peptide sequence. In this case the altered peptide sequence must be truncated to the first null amino acid, shortening the peptide. The total number of coalition vectors is determined by the product of the size of the background dataset and a predefined sampling parameter. For this work we set the background dataset size to 100 and the sampling parameter to 200. If a peptide was longer than 20, we increased this sampling parameter to 1000 for a more accurate estimate of the SV.

One consequence of substituting amino acids from the background dataset based on their N-C absolute positions are artifacts in the C-termini of peptides. The effect is most evident in the heatmaps of b-ions, for example there exists a light positive bias for all amino acids in Figure 3C for b7+1, approximately in the positions −2 to −6. This is the result of frequent truncation of peptides on their C-terminus when background substitutions are from shorter peptides than the one being explained. The particular effect, whether a negative or positive bias, depends on the model, and the position of the effect depends on the average length of peptides in the background dataset.

### Visualization

For analyzing the SV from all explained sequences in a validation set, we consider 2 aspects of any token: the amino acid type and the position in the sequence (AAP). When visualizing the overall mean values for any AAP, we set a minimum number of occurrences (15) for it to be displayed in our heatmaps. This is because some AAPs occur very rarely and thus a global explanation of their SV has little value for analysis.

For the intensity model we redefine the coordinate system based on the ion being explained. This coordinate system is based, set to 0, at the inner terminus, or break point, of the ion fragment, i.e. C-terminus of b-ions and N-terminus of y-ions. Positive numbers increase inside the fragment away from the breakpoint, and negative numbers decrease (more negative) outside of the fragment. For the iRT and CCS explanations we set up a binned coordinate system based on the length of the peptide. This is in effect the relative position along the length of the peptide, going from 0 at the N-terminus to 1 at the C-terminus.

## Acknowledgements

This work was in part funded by an ERC Starting Grant (Grant No. 101077037; ORIGIN; Joel Lapin and Mathias Wilhelm).

